# ALV-J and REV synergistically activate a new oncogene of KIAA1199 via NF-κB and EGFR signaling regulated by miR-147

**DOI:** 10.1101/338244

**Authors:** Defang Zhou, Jingwen Xue, Pingping Zhuang, Xiyao Cui, Shuhai He, Shuai Su, Guihua Wang, Li Zhang, Chengui Li, Libo Huang, Yingli Shang, Yongxiu Yao, Venugopal Nair, Huangge Zhang, ziqiang cheng

**Affiliations:** College of Veterinary Medicine, Shandong Agricultural University, Tai’an, 271018, China; The Pirbright Institute & UK-China Centre of Excellence on Avian Disease Research, Pirbright, Ash Road, Guildford, Surrey GU24 0NF, United Kingdom; Hancock Street, Department of Microbiology & Immunology, James Brown Cancer Center, University of Louisville, Louisville, KY 40202, USA

## Abstract

The tumorigenesis is the result of the accumulation of multiple oncogenes and tumor suppressor genes changes. Co-infection of avian leucosis virus subgroup J (ALV-J) and reticuloendotheliosis virus (REV), as two oncogenic retroviruses, showed synergistic pathogenic effects characterized by enhanced tumor initiation and progression. The molecular mechanism underlying synergistic effects of ALV-J and REV on the neoplasia remains unclear. Here, we found co-infection of ALV-J and REV enhanced the ability of virus infection, increased viral life cycle, maintained cell survival and enhanced tumor formation. We combined the high-throughput proteomic readout with a large-scale miRNA screening to identify which molecules are involved in the synergism. Our results revealed co-infection of ALV-J and REV activated a latent oncogene of KIAA1199 and inhibited the expression of tumor suppressor miR-147. Further, enhanced KIAA1199, down-regulated miR-147, activated NF-κB and EGFR were demonstrated in co-infected tissues and tumor. Mechanistically, we showed ALV-J and REV synergistically enhanced KIAA1199 by activation of NF-κB and EGFR signalling pathway, and the suppression of tumor suppressor miR-147 was contributed to maintain the NF-κB/KIAA1199/EGFR pathway crosstalk by targeting the 3’UTR region sequences of NF-κB p50 and KIAA1199. Our results contributed to the understanding of the molecular mechanisms of viral synergistic tumorgenesis, which provided the evidence that suggested the synergistic actions of two retroviruses could result in activation of latent pro-oncogenes.

**Author summary:** The tumorigenesis is the result of the accumulation of multiple oncogenes and tumor suppressor genes changes. Co-infection with ALV-J and REV showed synergistic pathogenic effects characterized by enhanced tumor progression, however, the molecular mechanism on the neoplasia remains unclear. Our results revealed co-infection of ALV-J and REV promotes tumorigenesis by both induction of a latent oncogene of KIAA1199 and suppression of the expression of tumor suppressor miR-147. Mechanistic studies revealed that ALV-J and REV synergistically enhance KIAA1199 by activation of NF-κB and EGFR signalling pathway, and the suppression of tumor suppressor miR-147 was contributed to maintain the NF-κB/KIAA1199/EGFR pathway crosstalk by targeting the 3’UTR region sequences of NF-κB p50 and KIAA1199. These results provided the evidence that suggested the synergistic actions of two retroviruses could result in activation of latent pro-oncogenes, indicating the potential preventive target and predictive factor for ALV-J and REV induced tumorigenesis.

## Introduction

Viral synergism occurs commonly in the nature when co-infection of two or more unrelated viruses invades the same host. As two oncogenic retroviruses, avian leukosis virus subgroup J (ALV-J) and reticuloendotheliosis virus (REV) are the optimal model to study the synergistic tumorigenesis mechanisms. Both ALV-J and REV consist of a set of retroviral genes, gag, pol, env and LTR, and mainly induce myelocytomas and reticuloendotheliosis, respectively (1, 2). Due to similar transmission routes, co-infection of ALV-J and REV can readily occur (3, 4), and spread very rapidly (5-7). Co-infection of ALV-J and REV caused more serious pathogenic effects, including growth retardation, immunosuppression, secondary infection and accelerated neoplasia progression in chickens (3, 5), leading to an increased mortality. Although the significance of the viral synergism had aroused high concerns, the synergistic tumorigenesis mechanisms, especially in retrovirus co-infections, remains unknown.

In synergistic interactions, biological traits such as virus accumulation, phenotypic and cytopathological changes, tissue tropism, host range and transmission rates of one or both of the viruses are changed (8, 9). Viruses may interact directly by transcomplementation of defective functions or indirectly, through host responses such as the defence mechanism (10-12). Initially, single viral infection was thought to be fully responsible for the tumorigenesis of each virus, while it is now established that in many cases, multiple viruses collaborate as co-factors in tumor formation. For example, viruses such as the Epstein Barr virus (EBV), the Kaposi’s sarcoma herpesvirus (KSHV), human immunodeficiency virus type 1 (HIV-1), human hepatitis C virus (HCV) show association with non-Hodgkin’s lymphomas (13). Co-infection of any one or multiple viruses may establish an environment that enhances tumor initiation and progression (14, 15). In additional to clinical phenotype, experimental models of co-infection have identified a variety of mechanisms that might contribute to tumorigenesis, including viral cofactors (16-18), common signalling pathway targets (18, 19), epigenetic modifications (reprogramming) (20-23), microenvironmental abnormalities (13) and interference with cell death (24, 25). However, whether the synergistic actions of two retroviruses result in activation of latent pro-oncogenes remains unclear.

For the retroviral genome packaging, ALV-J or REV needs to incorporate into equivalent amounts of cellular RNAs in vitro, which may direct or indirect active a latent oncogene (26-33). Most oncogenes have predominant roles in cancer, which include the receptor tyrosine kinase epidermal growth factor receptor (EGFR), nuclear factor-κB (NF-κB), the transcriptional regulator MYC, the small GTPase RAS and the phosphoinositide 3-kinase PI3K (34-39). Particularly, EGFR and NF-κB cooperate to provide oncogenic signals and promote tumor development by sustaining tumor cell proliferation, survival and invasiveness. NF-κB is activated upon EGFR, as well as NF-κB-dependent pathways underlying resistant to EGFR inhibitors (40). KIAA1199, which is reported in the Human Unidentified Gene-Encoded Large Proteins database, is a novel endoplasmic reticulum protein in cancer cell migration (41). In breast cancer, KIAA1199, as a NF-κB target gene, feeds back to impact on EGFR-dependent signalling.

MicroRNAs (miRNAs) constitute a large family of small noncoding RNAs functioning as major regulators of gene expressions in cancer development (42, 43). In consistent with single infection of retrovirus, co-infection of retrovirus and other virus also regulates the miRNA to mediate the cell signaling pathway through targeting the proto-oncogene. For instance, it has been verified that miR-718 and miR-891a-5p take part in regulating the PTEN/AKT/mTOR signaling pathway and NF-κB signaling pathway in the synergistic infection of HIV-1 and KSHV, respectively (18,44). MiR-147, upregulated by multiple TLRs in inflammatory response (45), had been reported to play a part in suppressing tumor by targeting EGFR-driven cell-cycle network proteins and inhibit cell-cycle progression and proliferation in breast cancer (46). Since a miRNA has multiple target genes, the molecular targets for miR-147 mediated regulation of inhibition of ALV-J and REV induced tumorigenesis remains to be identified.

Here, we investigated the synergistic mechanism of ALV-J and REV on tumorigenesis. We revealed that ALV-J and REV synergistically activate a latent oncogene KIAA1199 and inhibit a tumor suppressor miR-147. Furthermore, we demonstrated that the NF-κB/KIAA1199/EGFR pathway crosstalk that is targeted by miR-147 plays a pivotal role in the synergistic tumorigenesis of ALV-J and REV, providing the theoretical basis for further revealing the mechanism of tumor synergism.

## Results

### ALV-J synergizes REV to promote the viral replication and cell survival

To determine whether co-infection of ALV-J and REV has different effect on the recipient cells from single infection, we conducted viral RNA transcription and replication, protein expression and localization test. We caught the two viral particles of ALV-J and REV in the same cell by electron microscopy (Fig 1A), indicating co-infection of a single cell. The qRT-PCR results showed the RNA level of REV in co-infected cells was higher (P < 0.05) than those in single infected cells at 24 hpi to 96 hpi, and reached the highest peak at 72 hpi. ALV-J increased REV replication by 683.7-to 2882-fold (Fig 1B). In contrast, the RNA level of ALV-J showed no significant change between single and dual infection (Fig 1C). To further understand whether ALV-J and REV affect replication and subcellular localization each other in co-infected group, viral protein accumulation was observed dynamically by CLSM (Fig 1D). Dynamic analysis in single infected cells revealed that the ALV-J env protein accumulated in the cytoplasm and the nucleus at 48 hpi, and mainly localized in the cytoplasm with little remaining in the nucleus at 72 hpi. Accordingly, the REV env protein started accumulation in the cytoplasm of CEF at 72 hpi but not in the nucleus. Interestingly, in co-infected cells, the REV env protein started accumulation in the cytoplasm of cells at 48 hpi, and more ALV-J env protein production accumulated in cytoplasm than single infection. It is worth to note that ALV-J env protein started to appear in the nucleus at 96 hpi of co-infection. To further understand whether ALV-J synergizes REV to affect the proliferation rates of CEF cells, the number of the cells was tested by CCK-8 kit. Consequently, ALV-J and REV synergized with each other and further promote cell survival, indicating the high proliferation rate might associate with neoplasia (Fig 1E).

**Fig 1.**
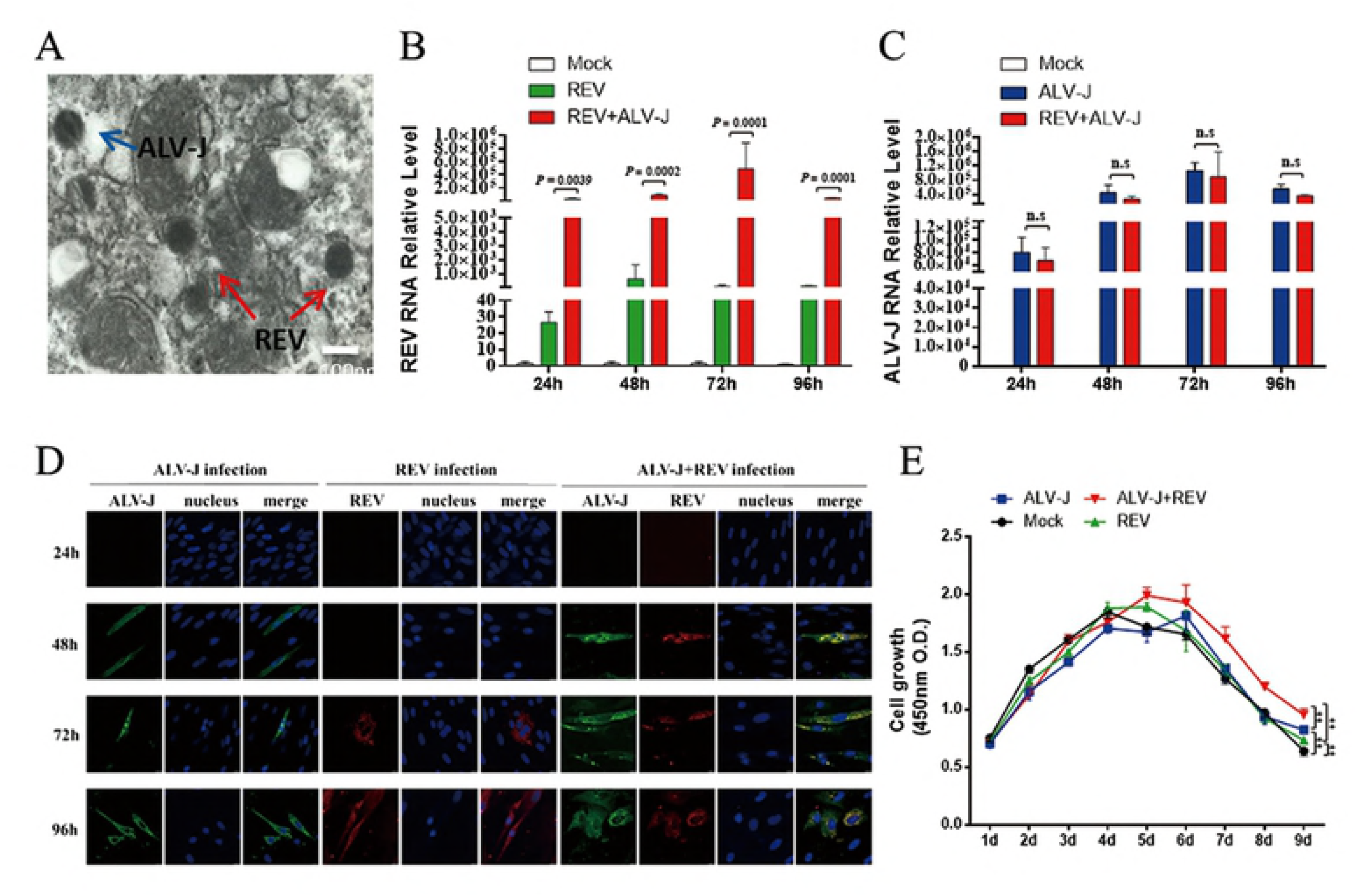
Co-infection of ALV-J and REV promotes the viral replication and cell survival in CEFs. (A) The two viral particles of ALV-J and REV were observed in the same cell by electron microscopy. (B) ALV-J increased REV RNA level at 24hpi, 48hpi, 72hpi and 96hpi. Data represent mean ± SEM determined from three independent experiments (n = 3), each experiment containing three technical replicates. (C) ALV-J RNA level was not changed significantly in intracellular co-infected ALV-J and REV at 24hpi, 48hpi, 72hpi and 96hpi. Data represent mean ± SEM determined from three independent experiments (n = 3), each experiment containing three technical replicates. (D) Co-infection of ALV-J and REV enhanced the viral proteins expression in intracellular. Protein expression of ALV-J(gag) and REV(env) by confocal in intracellular at 24hpi, 48hpi, 72hpi and 96hpi. (E) Co-infection of ALV-J and REV promoted CEFs survival. The CEFs were infected ALV-J, REV or both and further examined with CCK-8 assay on day 1, 2, 3, 4, 5, 6, 7,8 and 9 pi. Data represent mean ± SEM determined from three independent experiments (n = 6), each experiment containing three technical replicates.

### ALV-J synergizes REV to enhance pathogenicity and tumorigenesis

We established the animal model of co-inoculation of ALV-J and REV in allantoic cavity of fertilized SPF eggs at 6-day of embryonic age. The hatching rates, weights, H&E staining and qRT-PCR were used for confirming the synergistic pathogenicity and tumorigenesis of ALV-J and REV in vivo. The number of death in embryos showed that ALV-J synergized with REV to further decrease the hatching rate (Table 2). After hatching, both ALV-J and REV synergized with each other and further decreased the weights of chicken (Fig 2A). H&E staining results showed there are massive inflammatory cells that infiltrated around the blood vessel in liver and pancreas of chicken infected ALV-J, REV or both. Some lymphocytic focal inflammatory infiltrates were also observed in substantial areas of liver and kidney, and the demarcation between medulla and cortex in the thymus seemed to disappear (Fig 2B). The qRT-PCR results showed both viral RNA levels of liver, kidney, bursa of Fabricius, bone marrow and heart in co-infection group were higher (P < 0.05) than those in single infection group. REV increased ALV-J by 5.56-, 1.25-, 4.923-, 2.33-and 5.85-fold in liver, kidney, bone marrow, bursa of Fabricius and heart of co-infection group (Fig 2C), respectively, and ALV-J increased REV by 5.62-, 1.52-, 3.23-, 3.07-and 2.37-fold in the corresponding organs (Fig 2D), respectively. Further, some additional white nodules were observed in endocardium of co-infection group (Fig 2E). H&E staining results showed these white endocardial nodules were fibroma (Fig 2E). Collectively, co-infection of ALV-J and REV leads to induction of inflammation and tumorigenesis.

**Table 1.**
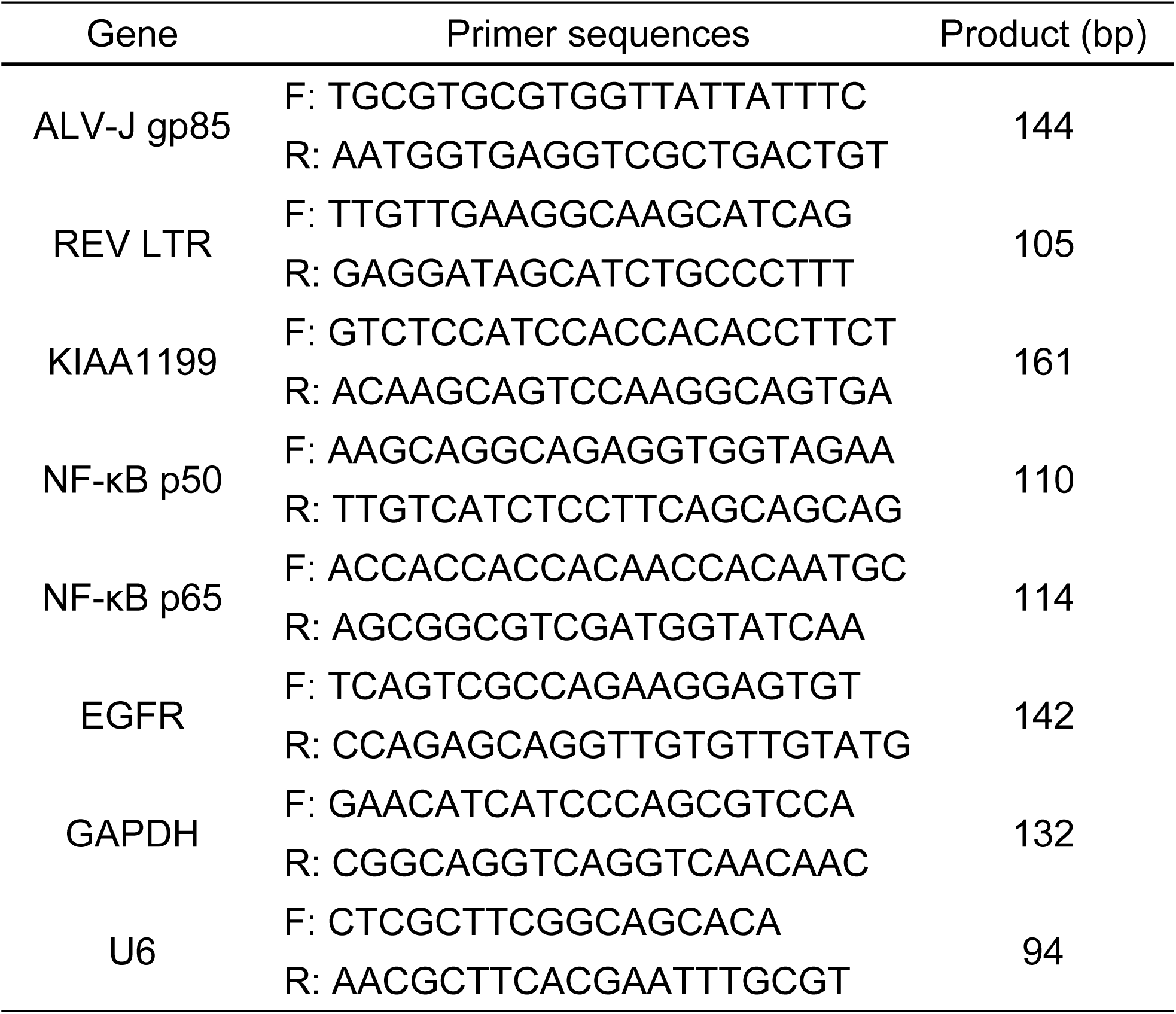
The primer sequences of real-time PCR.

**Table 2.**
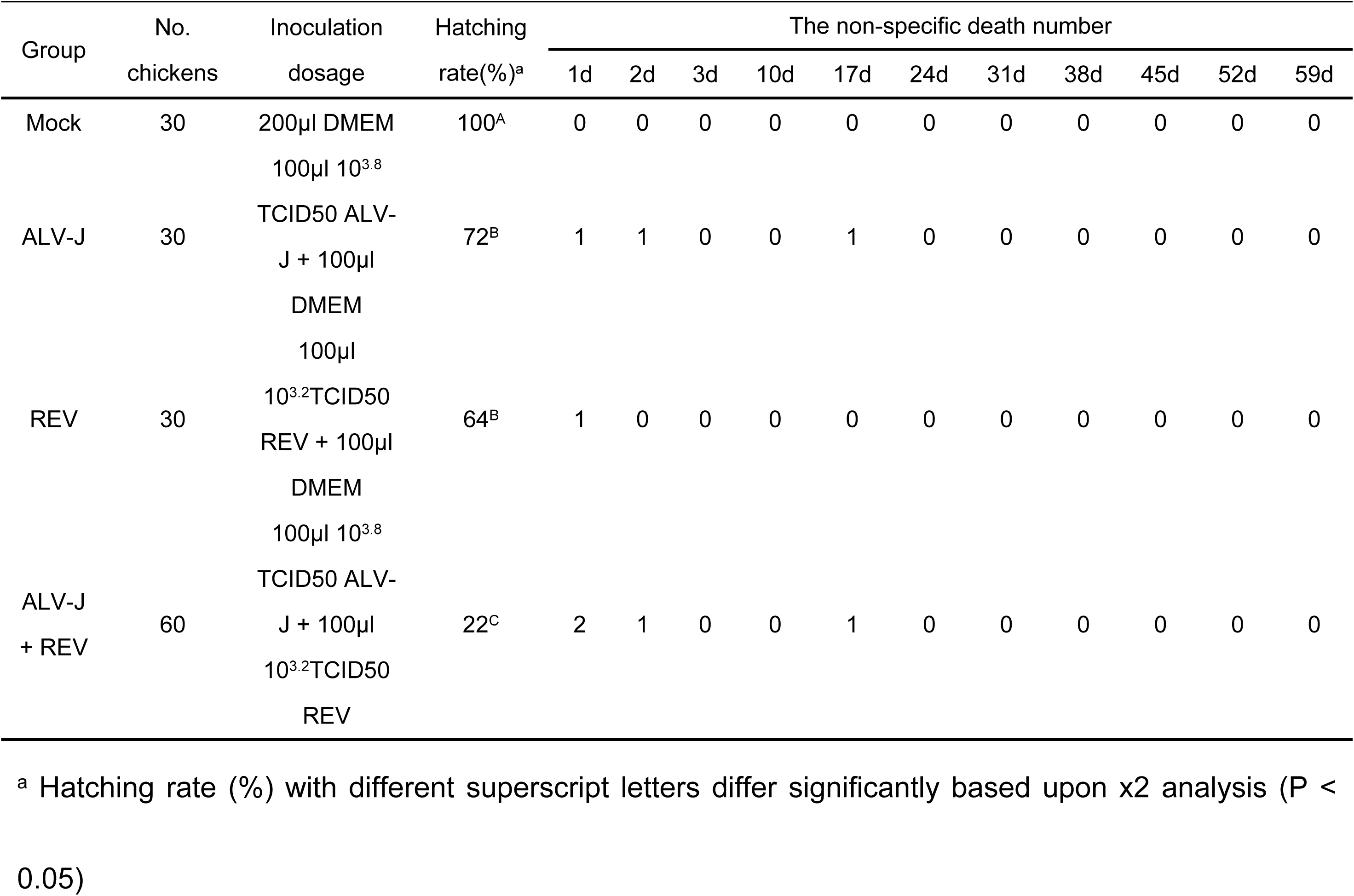
Influence of REV and ALV-J single infection and co-infection on the non-specific death number in chickens.

**Fig 2.**
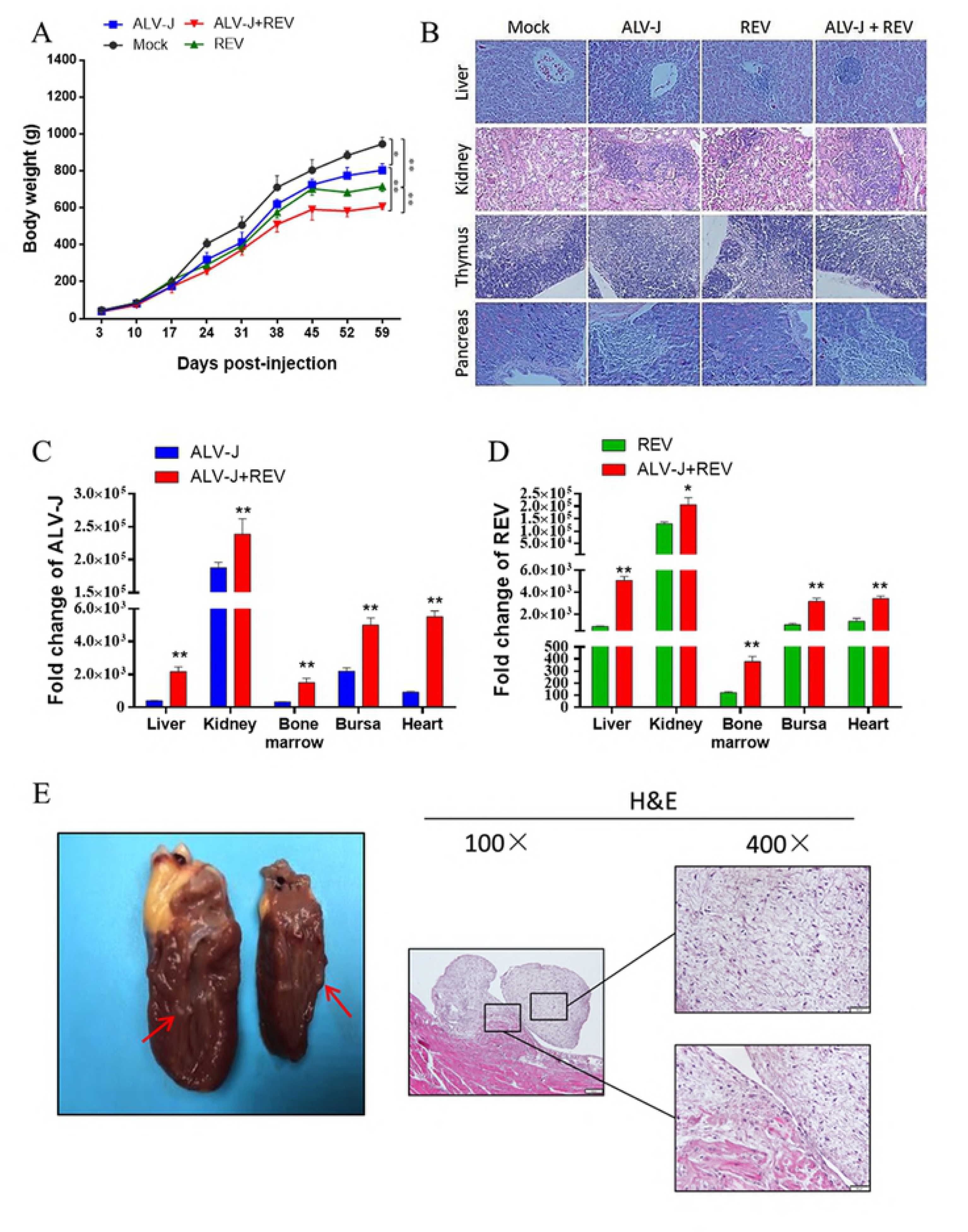
Co-infection of ALV-J and REV promoted the pathogenicity and tumorigenesis in chickens. (A) ALV-J synergized REV to decrease the weights of co-infected chickens. (B) ALV-J and REV synergistically promoted the pathogenicity in livers, kidneys, thymus and pancreas of chickens at 60 dpi. (C) REV increased ALV-J RNA level in liver, kidney, bone marrow, bursa of Fabricius and heart of chickens at 60 dpi. Data represent mean ± SEM determined from three independent experiments (n = 6), each experiment containing three technical replicates. Compared with single-infection group: *P < 0.05 and **P < 0.01. (D) ALV-J enhanced REV RNA level in liver, kidney, bone marrow, bursa of Fabricius and heart of chickens at 60 dpi. Data represent mean ± SEM determined from three independent experiments (n = 6), each experiment containing three technical replicates. Compared with single-infection group: *P < 0.05 and **P < 0.01. (E) Some additional white nodules were observed in endocardium of co-infection group. H&E staining results showed these white endocardial nodules were fibroma.

### ALV-J synergizes with REV to activate KIAA1199 and inhibit miR-147

To explore the synergistic tumorigenesis mechanism induced by ALV-J and REV, CEFs infected with ALV-J, REV or both were analyzed using iTRAQ quantitative proteomic analysis and miRNA whole-genome sequencing using the same batch of cell samples at 72 hpi. The protein thermodynamic chart showed there are 33 differentially expressed proteins in co-infected cells (Fig 3A). Interestingly, among the different expression proteins, KIAA1199 was declined in single-infected cells, while increased in co-infected cells. Further, western blot analysis showed ALV-J or REV decreased the KIAA1199 protein expression by 0.85-and 0.74-fold, respectively, while ALV-J synergized with REV to increase the KIAA1199 protein expression by 1.38-fold (Fig 3B). However, qRT-PCR result showed ALV-J, REV or both increased the KIA1199 RNA level by 2.59-, 2.41-and 3.89-fold, respectively (Fig 3C). The miRNA whole-genome sequencing showed miRNAs expression in the cells infected with ALV-J, REV, or both changed significantly compared to the Mock group. Interestingly, MiR-147 was the only downregulated miRNA in co-infection group, and we hypothesized that it has putative targeting sites in the 3’UTR of KIAA1199 (Fig 3D). The qRT-PCR result confirmed that co-infecting ALV-J and REV indeed decreased the expression of miR-147 in CEF cells, but CEF cell infected with single ALV-J or REV infection has increased the level of miR-147 (Fig 3E). To explore whether the above in vitro results were consistent with the chicken co-infected with ALV-J and REV at 60 dpi, we detected the RNA levels of KIAA1199 and miR-147 by qRT-PCR and protein expressions of KIAA1199 by IHC in liver, kidney, bone marrow, bursa of Fabricius and heart. The qRT-PCR results showed ALV-J synergized with REV to increase the KIAA1199 RNA level in liver, kidney, bone marrow bursa of Fabricius and heart (Fig 4A). Interesting, the IHC showed the expression levels of KIAA1199 were significantly increased in bone marrow, kidney and heart of ALV-J and REV co-infected group (Fig 4B and C), while that was not significantly increased in liver and bursa of Fabricius. The miR-147 expression was inhibited by co-infection of ALV-J and REV in kidney, bone marrow and heart, while that was increased in liver and bursa of Fabricius (Fig 4E), which was reversely co-related with KIAA1199 protein levels. Further, the high expression of KIAA1199 was also verified in endocardial fibroma by IHC. As expected, the level of KIAA1199 was significantly increased in fibroma cells and stroma and myocardial cells (Fig 4D), and the expression of miR-147 was decreased by ALV-J and REV co-infection (Fig 4E). In summary the data generated from both in vitro and in vivo demonstrated that co-infection of ALV-J with REV results in activation of KIAA1199 and inhibition of expression of miR-147.

**Fig 3.**
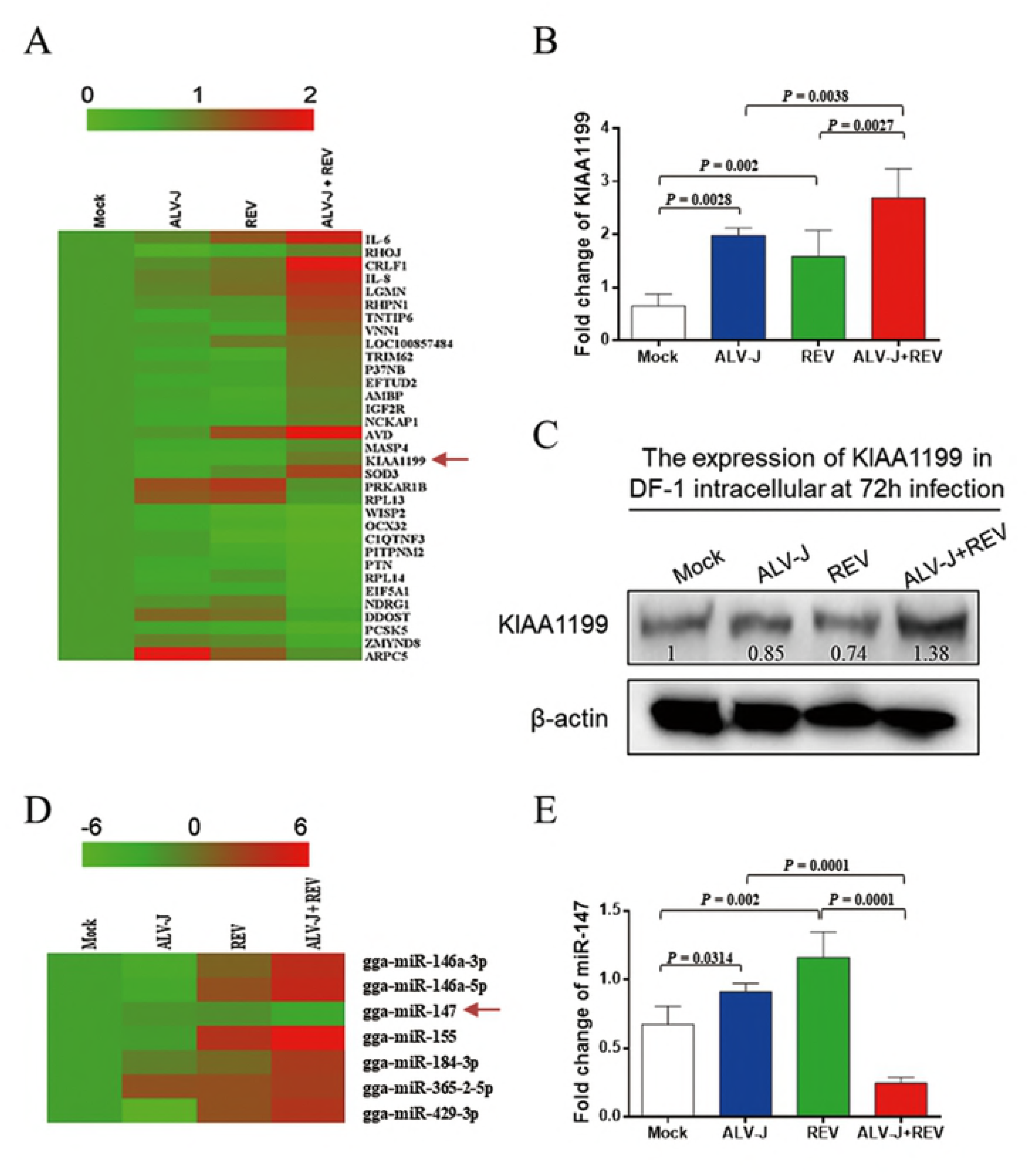
ALV-J synergized with REV to activate KIAA1199 and inhibit miR-147 in CEFs. (A) The iTRAQ quantitative proteomic analysis of infected CEF at 72hpi. There were 33 differentially expressed proteins between co-infection and single infection. (B) ALV-J and REV synergistically enhanced the KIAA1199 RNA level in infected DF-1 cells at 72hpi. Data represent mean ± SEM determined from three independent experiments (n = 3), each experiment containing three technical replicates. (C) ALV-J synergized with REV to enhance the KIAA1199 protein level in DF-1 cells at 72hpi detected by western blot with anti-KIAA1199 antibody. (D) MiR-147 was the only downregulated miRNA in co-infection group. (E) ALV-J synergized with REV to inhibit the miR-147 RNA level in DF-1 cells at 72hpi. Data represent mean ± SEM determined from three independent experiments (n = 3), each experiment containing three technical replicates.

**Fig 4.**
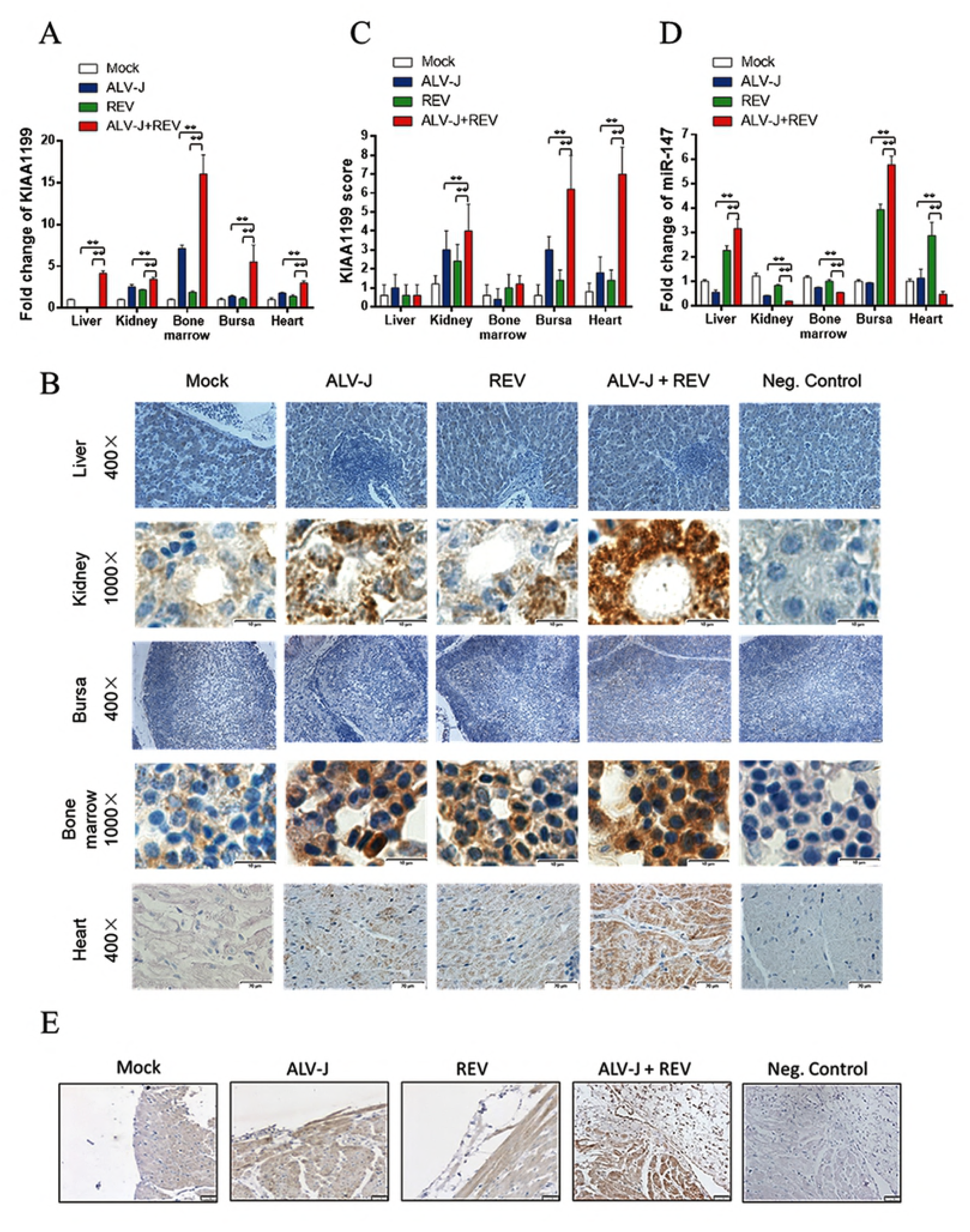
ALV-J synergized with REV to activate KIAA1199 and inhibit miR-147 in infected chicken. (A) ALV-J and REV synergistically enhanced the KIAA1199 RNA level in liver, kidney, bone marrow, bursa of Fabricius and heart of chicken at 60 dpi. Data represent mean ± SEM determined from three independent experiments (n = 6), each experiment containing three technical replicates. Compared with single-infection group: *P < 0.05 and **P < 0.01. (B) ALV-J synergized with REV to enhance the protein expression of KIAA1199 in kidney, heart and bone marrow in chicken at 60 dpi detected by immunohistochemical analysis with anti-KIAA1199 antibody. (C) Quantification of the results in (B). Compared with single-infection group: *P < 0.05 and **P < 0.01. (D) Co-infection of ALV-J and REV promoted the expression of KIAA1199 in fibroma of endocardium detected by immunohistochemical analysis with anti-KIAA1199 antibody. (E) The miR-147 expression was inhibited by co-infection of ALV-J and REV in kidney, bone marrow and heart, while that was increased in liver and bursa of Fabricius in chicken at 60 dpi. Data represent mean ± SEM determined from three independent experiments (n = 6), each experiment containing three technical replicates. Compared with single-infection group: *P < 0.05 and **P < 0.01.

### ALV-J and REV synergistically activated KIAA1199 via NF-κB and EGFR signaling

Previous study showed KIAA1199, which is transcriptionally induced by NF-κB proteins, promotes EGFR stability and signaling in breast cancer (40, 47). To explore the mechanism of KIAA1199 activation in mediating the synergistic effect of ALV-J and REV, the expression of NF-κB p65, p50 and EGFR were detected by qRT-PCR, western blotting and ELISA. The qRT-PCR of NF-κB results showed, compared with single infected ALV-J and REV, co-infection of ALV-J and REV increased the p65 RNA level by 1.78-and 1.91-fold, respectively (Fig 5A) and raised p50 RNA level by 2.51-and 2.03-fold, respectively (Fig 5B). The changes of western blot or ELISA results were consistent with above results that ALV-J synergizes with REV to further activate NF-κB signaling (Fig 5C and D). The qPCR of EGFR results also showed that ALV-J or REV decreased the EGFR RNA level by 0.42-and 0.81-fold, respectively, while ALV-J synergized with REV to increase the EGFR RNA level by 1.96-fold (Fig 5E). Further, western blot or ELISA results were consistent with the KIAA1199 protein expression in DF-1 cells infected with ALV-J, REV, or both. To explore whether the above results were consistent with the in vivo data of chicken infected with ALV-J and REV, the activation of NF-κB and EGFR RNA were detected using the chicken NF-κB P-IκBα ELISA kit and qRT-PCR in chicken co-infected with ALV-J and REV. The P-IκBα ELISA results showed that co-infection of ALV-J and REV increased the levels of phosphorylated IκBα in liver, kidney, bone marrow, bursa of Fabricius and heart (Fig 5F). Furthermore, ALV-J synergized with REV to enhance the EGFR levels in kidney, bone marrow, bursa of Fabricius and heart, while that decreased in liver (Fig 5G).

**Fig 5.**
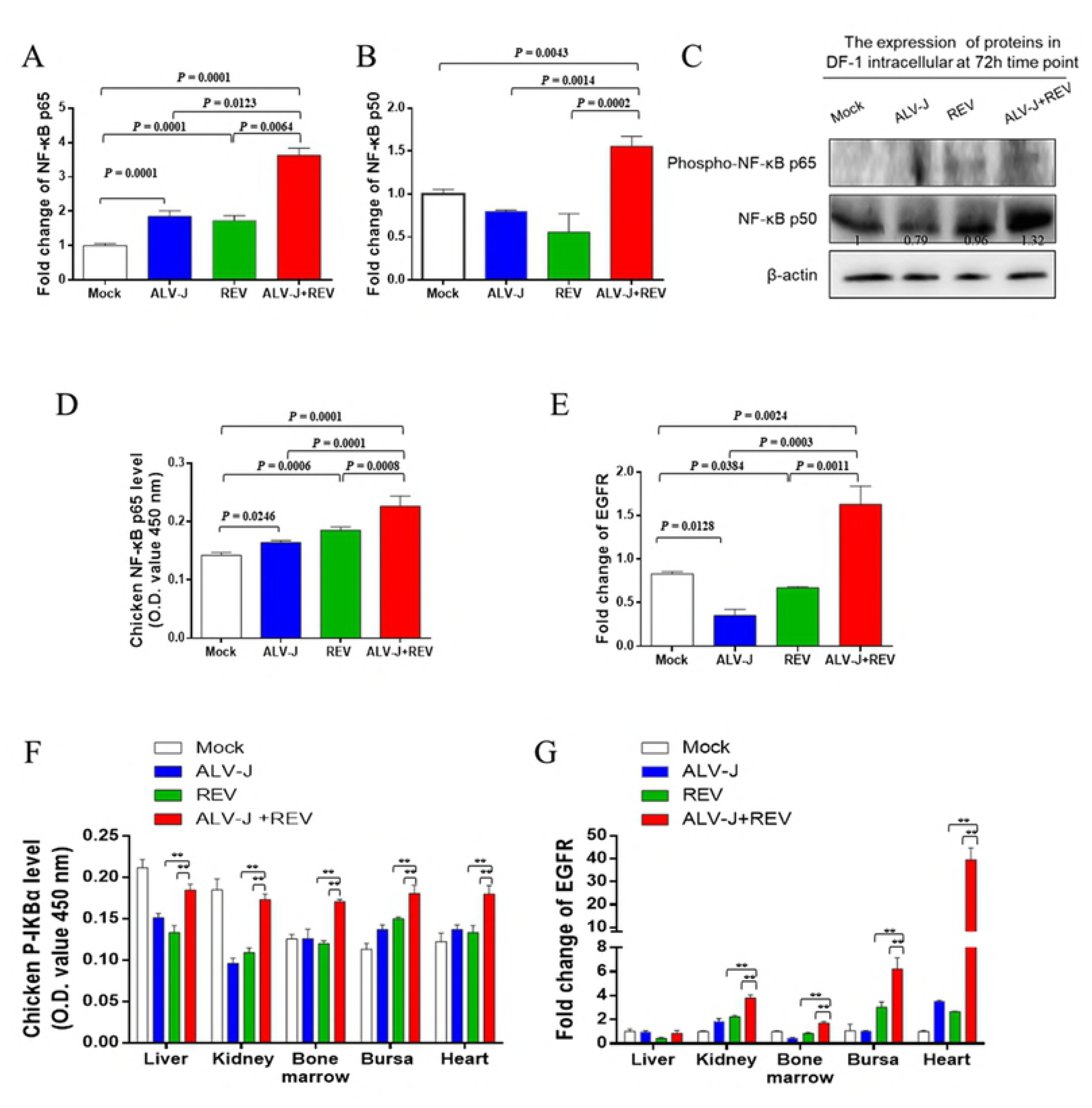
ALV-J and REV synergistically activated NF-κB and EGFR signaling. (A) Co-infection of ALV-J and REV increased the NF-κB p65 RNA level in DF-1 cells at 72hpi. Data represent mean ± SEM determined from three independent experiments (n = 3), each experiment containing three technical replicates. (B) ALV-J synergized with REV to promote the NF-κB p50 RNA level in DF-1 cells at 72 hpi. Data represent mean ± SEM determined from three independent experiments (n = 3), each experiment containing three technical replicates. (C) ALV-J synergized with REV to enhance the NF-κB p50 and phosphor-NF-κB p65 expression in DF-1 cells at 72hpi detected by western blot with anti-p50 and anti-phospho-p65 antibody. (D) ALV-J and REV synergistically enhanced the chicken NF-κB p65 protein level in DF-1 cells at 72hpi detected by Chicken NF-κB p65 ELISA kit. Competitive oligonucleotide was used as a positive control. Data represent mean ± SEM determined from three independent experiments (n=3), each experiment containing three technical replicates. (E) ALV-J synergized with REV to enhance EGFR RNA level in DF-1 cells at 72hpi. Data represent mean ± SEM determined from three independent experiments (n=3), each experiment containing three technical replicates. (F) ALV-J and REV synergistically enhanced the chicken P-IκBα expression in liver, kidney, bone marrow, bursa of Fabricius and heart in chicken at 60 dpi detected by Chicken NF-κB P-IκBα ELISA kit. Data represent mean ± SEM determined from three independent experiments (n=6), each experiment containing three technical replicates. Compared with single-infection group: *P < 0.05 and **P < 0.01. (G) ALV-J synergized with REV to increase the EGFR RNA level in liver, kidney, bone marrow, bursa of Fabricius and heart in chicken at 60 dpi. Data represent mean ± SEM determined from three independent experiments (n=6), each experiment containing three technical replicates. Compared with single-infection group: *P < 0.05 and **P < 0.01.

To further determine whether NF-κB is required for induction of KIAA1199 and EGFR, we constructed and transfected the NF-κB p65 shRNAs into the DF-1 cells. Indeed, when NF-κB p65 was knockdown, the RNA levels of KIAA1199 and EGFR were decreased (Fig 6A, B and C). Identitical results were observed in KIAA1199 inhibition (Fig 6D, E and F).

**Fig 6.**
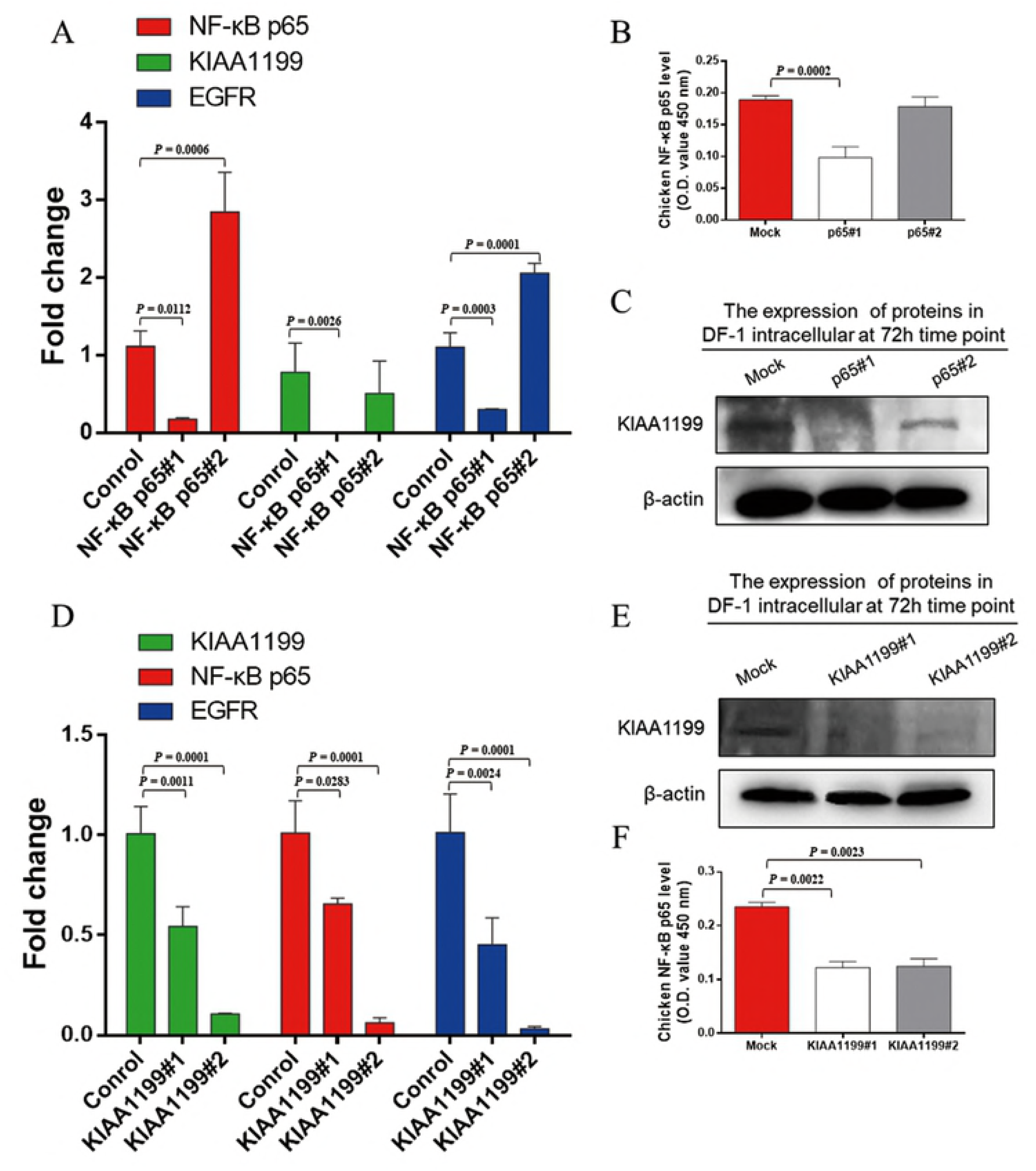
The KIAA1199 was activated by NF-κB and EGFR signaling during ALV-J and REV infection. (A) The RNA levels of NF-κB p65, KIAA1199 and EGFR were suppressed by incubating NF-κB p65 shRNA at 48hpi. Data represent mean ± SEM determined from three independent experiments (n=3), each experiment containing three technical replicates. (B) The chicken NF-κB p65 protein level was inhibited by NF-κB p65 shRNA detected by Chicken NF-κB p65 ELISA kit. Data represent mean ± SEM determined from three independent experiments (n = 3), each experiment containing three technical replicates. (C) The protein expression of KIAA1199 was decreased by NF-κB p65 shRNA at 48hpt detected by western blot with anti-KIAA1199 antibody. (D) The RNA level of KIAA1199, NF-κB p65 and EGFR were suppressed by KIAA1199 shRNA. Data represent mean ± SEM determined from three independent experiments (n = 3), each experiment containing three technical replicates. (E) The protein expression of KIAA1199 was decreased by KIAA1199 shRNA at 48h pt detected by western blot with anti-KIAA1199 antibody. (F) The protein level of chicken NF-κB p65 was inhibited by KIAA1199 shRNA detected by Chicken NF-κB p65 ELISA kit. Data represent mean ± SEM determined from three independent experiments (n = 3), each experiment containing three technical replicates.

Together these data suggest that ALV-J and REV synergistically activate the NF-κB /KIAA1199/EGFR pathway.

### Cellular miR-147 targets NF-κB /KIAA1199/EGFR pathway

To explore whether miR-147 directly target KIAA1199, the relationship of miR-147 and KIAA1199 were verified by dual-luciferase assay. Indeed, KIAA1199 3’UTR luciferase reporter assay showed that miR-147 significantly inhibited the activity of KIAA1199 3’UTR reporter but not that of the control reporter (Fig 7A). MiR-147 inhibited the activity of the KIAA1199 3’UTR reporter and the endogenous KIAA1199 expression in CEF in a dose-dependent manner (Fig 7B and 7C). On the contrary, a miR-147 inhibitor up-regulated the endogenous KIAA1199 expression level in a dose-dependent manner (Fig 7D). Bioinformatics analysis identified one putative miR-147 binding site at the KIAA1199 3’UTR (Fig 7E). Mutation of this site abolished the miR-147 inhibitory effect on KIAA1199 3’UTR reporter activity (Fig 7F). These data suggest that miR-147 directly targets KIAA1199.

**Fig 7.**
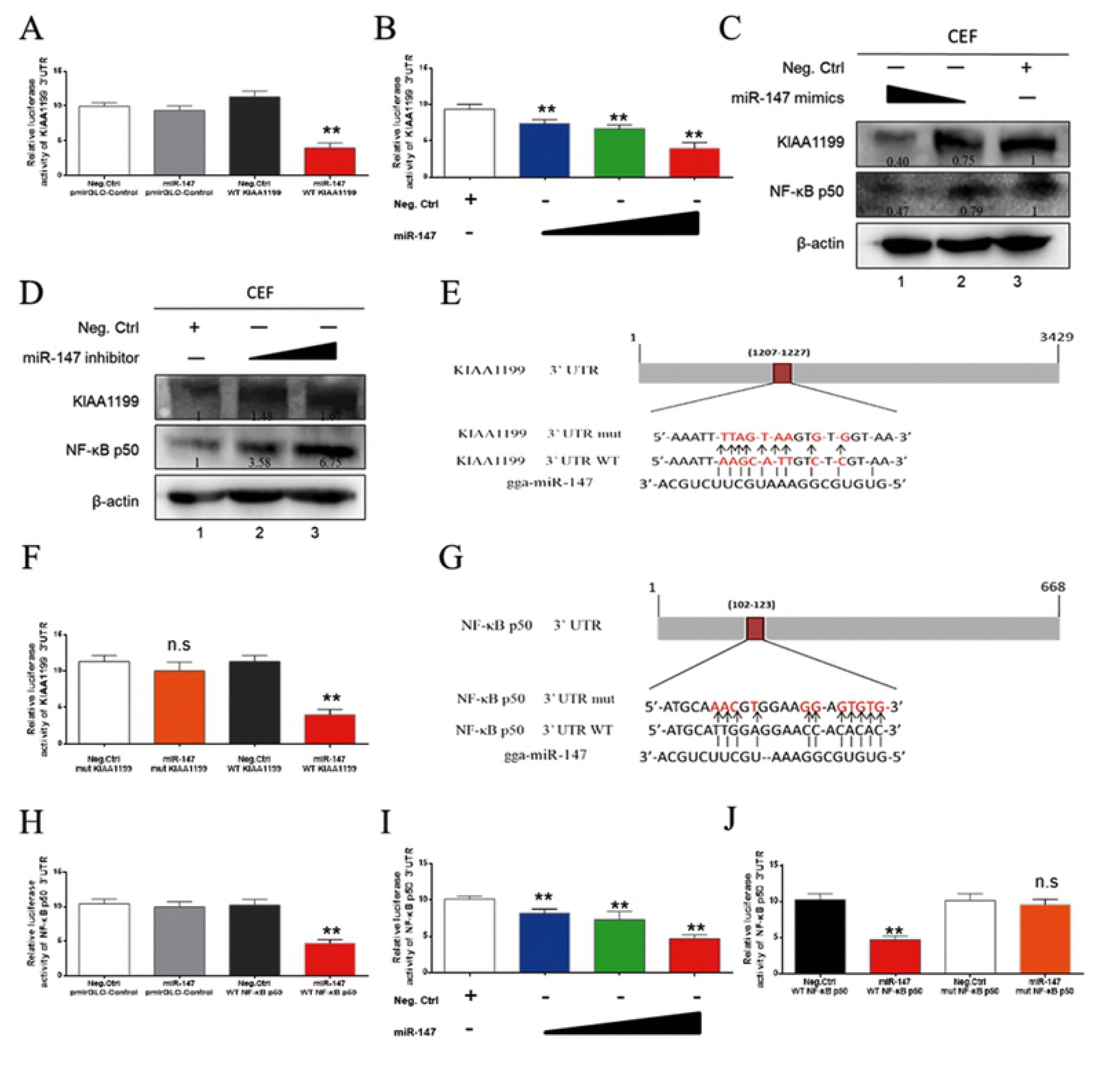
Cellular miR-147 regulated NF-κB p50 and KIAA1199 by targeting 3’UTR. (A) MiR-147 inhibited the reporter activity of the pmirGLO-KIAA1199 3’UTR. MiR-147 mimics (40 nM) or negative control was co-transfected with pmirGLO-Control or pmirGLO-KIAA1199 3’UTR reporter plasmid into 293T cells. (B) MiR-147 mimics (10, 20 and 40 nM) or a negative control was co-transfected along with pmir-GL0-KIAA1199 3’UTR reporter plasmid into 293T cells. (C) MiR-147 inhibited expressions of KIAA1199 and NF-κB p50 in a dose-dependent manner. MiR-147 mimics (20 and 40 nM) were transfected into ALV-J and REV co-infected CEF, western blot was performed with anti-KIAA1199 antibody or anti-NF-κB p50 antibody at 48 h post-transfection. (D) Inhibition of miR-147 promoted expression of KIAA1199 and NF-κB p50. MiR-147 inhibitors (30 and 60 nM) were transfected into CEF and western blotting was performed with anti-KIAA1199 or anti-NF-κB p50 antibody at 48 h post-transfection. (E) Schematic diagram of predicted seed sequence of miR-147 which binds with KIAA1199 3’UTR. (F) KIAA1199 3’UTR wild type (WT KIAA1199) was co-transfected with a negative control (Neg. Ctrl.) or miR-147 into 293T cells, while mutant KIAA1199 3’UTR construct (mut KIAA1199) was also co-transfected with Neg. Ctrl. or miR-147. (G) Schematic diagram of predicted seed sequence of miR-147 which binds with NF-κB p50 3’UTR. (H) MiR-147 inhibited the reporter activity of the pmirGLO-NF-κB p50 3’UTR. MiR-147 mimics (40 nM) or negative control with pmirGLO-Control or pmirGLO-NF-κB p50 3’UTR reporter plasmid were cotransfected into 293T cells. (I) MiR-147 mimics (10, 20 and 40 nM) or a negative control was co-transfected into 293T cells along with pmir-GL0-NF-κB p50 3’UTR reporter plasmid. (J) NF-κB p50 3’UTR wild type (WT KIAA1199) was co-transfected with a negative control (Neg. Ctrl.) or miR-147 into 293T cells, while mutant NF-κB p50 3’UTR construct (mut NF-κB p50) was also co-transfected with Neg. Ctrl. or miR-147. All above Luciferase assays were performed 48 h later, data represent the mean ± SEM from three independent experiments (n = 3), and each experiment containing three technical replicates. **P < 0.01 by Student’s t test versus the Neg. Ctrl. group. n.s., not signifcant.

Interestingly, previous study showed miR-147 is a member of suppressors for targeting EGFR signaling proteins (46). As the close cooperation of NF-κB, KIAA1199 and EGFR, miR-147 may also have some connection to NF-κB in mediating the synergistic effect of ALV-J and REV. To evaluate the relationship with NF-κB signaling, miR-147 was also predicted with the related proteins. As expected, bioinformatics analysis identified one putative miR-147 binding site in the 3’UTR of NF-κB p50 (Fig 7G). Further, NF-κB p50 3’UTR luciferase reporter assay showed that miR-147significantly inhibited the activity of NF-κB p50 3’UTR reporter but not that of the control reporter (Fig 7H). MiR-147 inhibited the activity of the NF-κB 3’UTR reporter and the NF-κB p50 expression in CEF in a dose-dependent fashion (Fig 7C and I). A miR-147 inhibitor up-regulated the NF-κB p50 expression level in a dose-dependent manner (Fig 7D). Mutation of this site abolished the miR-147 inhibitory effect on NF-κB p50 3’UTR reporter activity (Fig 7J). These data suggest that miR-147 also directly targets NF-κB p50 besides KIAA1199.

## Discussion

In this study, a specific synergism was observed in ALV-J and REV co-infected CEFs that characterized by extremely enhanced the ability of virus infection, increased viral life cycle, maintained cell survival, and chickens that characterized by severe weight loss, more shedding virus (viral loading), more serious pathological changes including severe inflammation, enhanced tumor formation and extended tumor spectrum. One of molecular mechanisms underlying the synergism is demonstrated in this study. We found that co-infection of ALV-J and REV activates NF-κB mediated inflammation pathway that causes upregulation of p50 and p65, leading to increased KIAA1199 gene transcription. KIAA1199 also activate EGFR signaling to promote NF-κB pathway stability, and finally formed the signaling pathway crosstalk. Furthermore, the suppression of tumor suppressor miR-147 was contributed to maintain the NF-κB/KIAA1199/EGFR pathway crosstalk by targeting the 3’UTR region sequences of NF-κB p50 and KIAA1199 (Fig 8).

**Fig 8.**
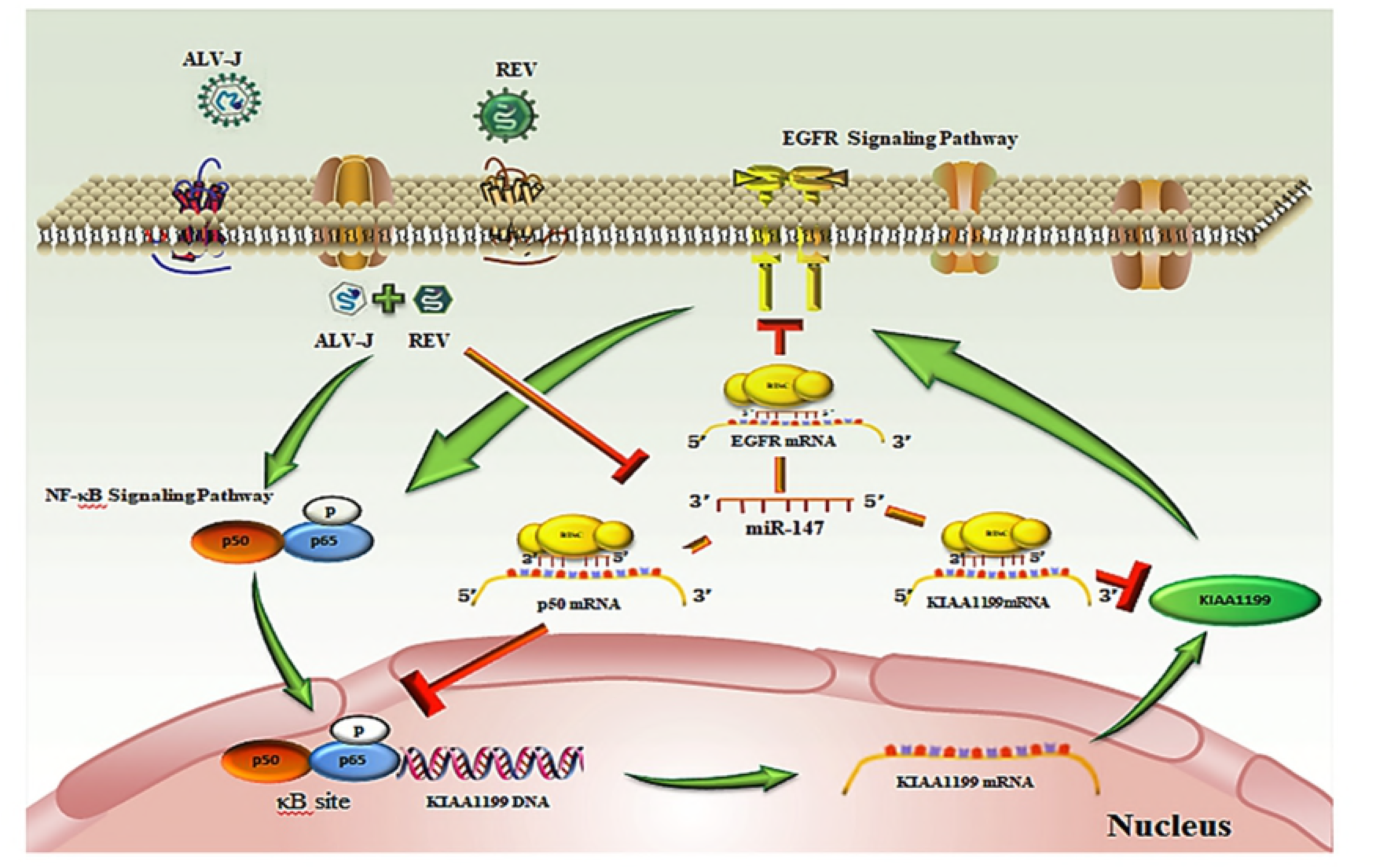
Diagram of molecular mechanism by which ALV-J and REV synergistically activate KIAA1199 and inhibit miR-147. In ALV-J and REV co-infected cells, ALV-J synergizes with REV to further promote the NF-κB pathway. As a result, p50 and p65 were enhanced, leading to drive KIAA1199 gene transcription. KIAA1199 also activate EGFR signaling to promote NF-κB pathway stability. MiR-147, as a key to NF-κB/KIAA1199/EGFR pathway crosstalk, regulated p50, KIAA1199 and EGFR-driven cell-cycle proteins by targeting their 3’UTR region sequences, respectively, to further active the oncogene KIAA1199.

Previous studies on the mechanisms of tumor initiation and progression by synergistic interactions had been limited in viral cofactors (16-18), common signalling pathway targets (18, 19), epigenetic modifications (reprogramming) (20-23), microenvironmental abnormalities (13) and interference with cell death (24, 25). In current study, we found ALV-J and REV synergistically activated a new oncogene of KIAA1199 by the inhibition of miR-147 and enhancement of NF-κB, which provided the powerful evidence that suggested the synergistic actions of two retroviruses could result in activation of latent pro-oncogenes.

Mechanistic studies revealed that ALV-J and REV synergistically enhances the expression of KIAA1199 via NF-κB mediated pathway. We also verified ALV-J synergizes with REV to promote NF-κB signaling and EGFR signaling in vitro and vivo. Further, the results of the NF-κB p65 and KIAA1199 shRNA tests validated the existence of NF-κB /KIAA1199/EGFR pathway crosstalk in co-infection of ALV-J and REV indirectly. However, the protein expression of KIAA1199 and its down-stream EGFR were inhibited in single infected ALV-J or REV. Therefore, the different regulations of KIAA1199 may provide an important role of tumor progress. Previous studies showed that as an oncogene, KIAA1199 expression is enhanced in breast, gastric and colon cancer (41, 48-51). In current study, ALV-J synergized with REV to increase the KIAA1199 expression in bone marrow, kidney, heart and endocardial fibroma, indicating the tissue specificity of KIAA1199 and the correlation to tumor spectrum extending.

Previous studies have shown that both NF-κB and EGFR signaling pathways could be activated by single-infecting ALV-J or REV. For instant, ALV-J induces VEGF expression via NF-κB/PI3K-dependent IL-6 production to promote tumorigenesis (52-54). In ALV-induced erythroblastosis, c-erbB encodes the carboxyl-terminal domain of the epidermal growth factor receptor (55, 56). NF-κB activation increased in expression of v-rel avian reticuloendotheliosis viral oncogene homolog (c-REL) to transduce and activate EGFR (57). As the considerable participators in various biological processes, NF-κB and EGFR pathways are involved in different stages of tumorigenesis induced by viruses, such as invasion, proliferation of tumor cells, apoptosis, migration and angiogenesis. While growth factors trigger NF-kB-activating cascades upon binding to ERBB members, the transcriptional induction of some NF-kB target genes, such as KIAA1199, also feeds back to impact on EGFR-dependent signaling pathways. Although this similar phenomenon had been verified in breast cancer or neck squamous cell carcinomas (HNSCCs) (47, 58-60), indicating this crosstalk was emerged as the results by viruses synergistic infection.

Compared to single infection, the protein expression of KIAA1199 was only increased in co-infection group but not single infection, while the KIAA1199 RNA levels was increased in each group, implying that co-infection of ALV-J with REV regulated KIAA1199 post transcriptionally, and there were also some key molecular regulators to determine the expression of KIAA1199 protein. Among many of the miRNA biomarkers, miR-147 was still recognized as an important factor in inflammatory responses and cancer. In small-cell lung cancer, miR-147 was found to be closely associated with cancer chemo resistance (61). MiR-147 is expressed in normal lung tissues and was upregulated as part of the inflammatory responses (45). In this study, the expressions of miR-147 were decreased in kidney, bone marrow and heart, which were correlated with the oncogene KIAA1199 activation by co-infection of ALV-J and REV. Conversely, the KIAA1199 levels were inhibited by miR-147 in liver and bursa of Fabricius suggesting that miR-147 also acted as a suppressor (62). MiR-147 was also a potential suppressor that targets EGFR-driven cell-cycle network protein in breast cancer (46). As NF-κB, KIAA1199 and EGFR pathway were connected closely in cancer, miR-147 was also analysed with NF-κB signaling proteins. As expected, bioinformatics analysis also identified one putative miR-147 binding site in the 3’UTR of NF-κB p50. Relative to the suppression for EGFR-driven cell-cycle protein, miR-147 was a key to NF-κB/KIAA1199/EGFR pathway crosstalk activation during ALV-J synergizes with REV. This initial finding will open up a new avenue for further identifying factors that regulate the stability of KIAA1199 protein.

In summary, our results demonstrate that co-infection of ALV-J and REV promotes tumorigenesis by both induction of a latent oncogene of KIAA1199 and suppression of the expression of tumor suppressor miR-147 through activation of NF-κB and EGFR signalling pathway crosstalk. As a negative feedback mechanism, KIAA1199 is also directly targeted by overexpressed miR-147 for inhibition of production of KIAA1199 protein. Our results contributed to the understanding of the molecular mechanisms of viral synergistic tumorgenesis, which provided the powerful evidence that suggested the synergistic actions of two retroviruses could result in activation of latent pro-oncogenes.

## Methods

### Cells, virus, and plasmids

Human 293T cells were purchased from Procell Life Science & Technology Ltd., China. Avian DF-1 cells and chicken embryo fibroblasts (CEFs) cells were propagated and maintained in Dulbecco’s Modified Eagle’s Medium (DMEM) supplemented with 10% fetal bovine serum (FBS), 1% penicillin/streptomycin, 1% l-glutamine, and in a 5% CO2 incubator at 37 ° C (63, 64).The stock SNV strain of REV at 10^3.2^ 50% tissue culture infectious doses (TCID_50_) and NX0101 strain of ALV-J at 10^3.8^ TCID_50_ were maintained in our laboratory, respectively. The TCID_50_ of SNV and NX0101 strain were titrated by limiting dilution in DF-1 culture. The KIAA1199 and NF-κB p50 3’UTR were cloned into the downstream of luciferase reporter gene of pmirGLO Control vector to create wild type pmirGLO-KIAA1199 3’UTR (WT KIAA1199) and pmirGLO-p50 3’UTR (WT p50) plasmids, respectively (GenePharma, Shanghai, China). Bioinformatics analysis software tools and web sites, including RNA22, NCBI and ENSEMBLE, were used to analyse and predict the binding sites between miR-147 and 3’UTR of KIAA1199 or NF-κB p50. The construction of the pmirGLO-KIAA1199 3’UTR mutant plasmid (Mut KIAA1199) and the pmirGLO-p50 3’UTR mutant plasmid (Mut p50) were constructed by site-directed mutagenesis.

### Pathogenicity and tumorigenesis assay in SPF chickens

Specific pathogen free (SPF) chicken embryos were purchased from the SPAFAS Co. (Jinan, China; a joint venture with Charles River Laboratory, Wilmington, MA, USA). 150 SPF chicken embryos were divided into 4 groups placed in separated incubators receiving filtered positive-pressure air. All embryos were inoculated via allantoic cavity at 6-day of embryonic age with ALV-J of 100μl 10^3.8^TCID_50_ and 100μl DMEM per egg (30 eggs), REV of 100μl 10^3.2^ TCID_50_ and 100μl DMEM per egg (30 eggs), or both ALV-J of 100μl 10^3.8^ TCID_50_ and REV of 100μl 10^3.2^ TCID_50_ per egg (60 eggs).The embryos from Mock were inoculated with 200μl DMEM (30 eggs). After hatching, the shedding virus was tested by ELISA for p27 antigen of ALV-J from cloacal swabs or qPCR for REV LTR gene from blood, respectively. All birds were weighed and clinical performance was recorded each week. To observe the pathogenicity and tumorigenesis, 6 chickens from each group were euthanatized and examined post-mortem at 60 days. Every tissue from each chicken was divided into four portions for histopathology, immunohistochemistry (IHC), ELISA and western blot assay. The RNA was extracted from liver, kidney, bone marrow, bursa of Fabricius and heart to detect the viral RNA, KIAA1199, EGFR or miR-147.

### iTRAQ-labelled LC-MS/MS

SDT buffer was added to the CEF sample either Mock or infected with ALV-J or REV alone, or co-infected with both ALV-J and REV for 72 hpi. The lysate was sonicated and then boiled for 15 minutes. After centrifuged at 14, 000×g for 40 minutes, the supernatant was quantified with the BCA Protein Assay Kit (Bio-Rad, USA). Proteins for each sample were separated by SDS-PAGE, prepared by sample filter-aid, and labelled using iTRAQ reagent according to the manufacturer’s instructions (Applied Biosystems). The iTRAQ labelled peptides were fractionated by SCX chromatography using the AKTA Purifier system (GE Healthcare). Each fraction was injected for nanoLC-MS/MS analysis. LC-MS/MS analysis was performed on a Q Exactive mass spectrometer (Thermo Scientific). MS/MS spectra were searched using MASCOT engine (Matrix Science, London, UK; version 2.2) embedded into Proteome Discoverer 1.4.

### Illumina small RNA deep sequencing

Total RNA of the infected CEF sample, the same batch of samples as in iTRAQ assay, was separated by 15% agarose gels to extract the small RNA (18-30 nt). After precipitated by ethanol and centrifugal enrichment of small RNA population, the library was prepared according to the method and process of Small RNA Sample Preparation Kit (Illumina, RS-200-0048). RNA concentration of the library was measured using Qubit^®^ RNA Assay Kit in Qubit^®^ 2.0 to preliminary quantify and then dilute to 1ng/µl. Insert size was assessed using the Agilent Bioanalyzer 2100 system (Agilent Technologies, CA, USA), The library with the expected insert size was then quantified accurately using Taqman fluorescence probe of AB Step One Plus Real-Time PCR system (Library valid concentration>2nM).The qualified libraries were sequenced by an Illumina Hiseq 2500 platform and 50 bp single-end reads were generated.

### RNA interference

Two shRNAs targeting each gene were co-transfected into DF-1. shRNAs, 5’-GCATTGAGGAGAAGCGCAAAC-3’ and 5’-GAAAGAGGACATCGAGGTTCG-3’ targeting NF-κB p65, and 5’-GCTGGAATGATCATCGATAAC-3’ and 5’-GCATGGAAGAAGGCGGATATT-3’ targeting KIAA1199 were purchased from GenePharma (Shanghai, China).

### MiRNA mimics and inhibitors

The sequence of commercially available miR-147 mimics was 5’-GUGUGCGGAAAUGCUUCUGCAGAAGCAUUUCCGCACACUU-3’; the negative control sequence was 5’-UUCUCCGAACGUGUCACGUTT-3’; 5’-ACGUGACACGUUCGGAGAATT-3’; the sequence of miR-147 inhibitor was 5’-GCAGAAGCAUUUCCGCACAC-3’; and the inhibitor negative control sequence was 5’-CAGUACUUUUGUGUAGUACAA-3’. MiRNA mimics and miRNA inhibitor were purchased from GenePharma (Shanghai, China).

### Luciferase reporter assay

The miRNA mimics, KIAA1199 3’UTR or NF-κB p50 3’UTR luciferase reporter plasmid were co-transfected into 293T cells. At 48hpi, cell lyases were prepared according to the manufacture’s instruction. Luciferase activity was measured with a Dual-Luciferase Reporter Assay System (Beyotime Co., Ltd) and normalized against the activity of the Renilla luciferase gene.

### Western blotting

To test KIAA1199, NF-κB p65 and p50 expression, DF-1 cells were lysed in cell lysis buffer (Beyotime) and incubated on ice for 5 minutes. The lysates were resuspended in SDS loading buffer, boiled for 5 minutes, loaded and run on a 6% (KIAA1199) and 10% (NF-κB p65 and p50) SDS-PAGE, respectively, and then transferred onto a nitrocellulose membrane (Solarbio). The nitrocellulose membrane was blocked with 5% skimmed milk at 4°C overnight and probed with anti-KIAA1199 (Bioss), anti-p50 (Abways) and anti-phospho-p65 (Abways) antibody at a 1:100, 1:500 and 1:5000 dilution, respectively, followed by horseradish peroxidase (HRP)-conjugated goat anti-mouse secondary antibody (Bioss) at a 1:3000 dilution. The beta-actin was used as loading control. Detection was performed with Enhanced HRP-DAB Chromogenic Substrate Kit (Tiangen) according to the manufacturer’s instructions.

### Transmission electron microscopy

The protocol was conducted as described in the previous study (65).

### Confocal microscopy assay

For confocal microscopy assay, the cells were washed three times with PBS (137mM NaCl, 2.7mM KCl, 10mM Na_2_HPO_4_, and 2mM KH_2_PO_4_) and were fixed in fixative (1ml of fixative contains 600μl of acetone plus 400μl of alcohol) at 24h, 48h, 72h and 96hpi. 7 minutes later, the fixative was removed and the cells were washed three times with PBS. Then, the cells were incubated with anti-env of REV (1:200) or anti-gp85 of ALV-J (1:200) antibody at 4°C for 10h. After that, the cells were washed three times with PBS, and incubated with HRP-labelled Goat anti-rabbit IgG (H+L) (1:5000) and HRP-labelled Goat anti-mouse IgG (H+L) (1:5000) at 37°C for 1.5h. After washing three with PBS, Nucleuses were stained with DAPI at 37°Cfor 5 minutes, and then the cells were washed three times with PBS again. Finally, the cells with 1ml of 50% glycerol were observed in Confocal laser scanning microscopy (CLSM) immediately.

### Hematoxylin and eosin (H&E) and immunohistochemical staining

The liver, kidney, bone marrow, pancreas, bursa of Fabricius and thymus of chicken were formalin-fixed, paraffin embedded, sectioned, and stained for histopathology observation. For immunohistochemical analysis, paraffin sections were deparaffinized and rehydrated in graded alcohols and antigen retrieval was performed using citrate buffer in a pressure cooker for 6 minutes. The home-made anti-KIAA1199 antibody (1:100) was used for the primary reaction. Immunoperoxidase staining was performed using the LSAB kit (Dako, Glostrup, Denmark), according to the supplier’s recommendations. Positive cells were visualized using a 3, 3’-diaminobenzidine substrate and the sections were counterstained with haematoxylin.

The immunolabelled tissues were evaluated by using a semi-quantitative score of the intensity and extent of the staining according to an arbitrary scale (47).

### Real-time quantitative reverse transcription polymerase chain reaction

The specific primer sequences of KIAA1199, NF-κB p50, NF-κB p65, EGFR and GAPDH were listed in Supplementary Table S1. Total RNA from CEF either Mock, single infection of ALV-J or REV, or co-infection of both ALV-J and REV, were isolated using Tiangen RNeasy mini kit as per manufacturer’s instructions, with optional on-column DNase digestion, respectively. RNA integrity and concentration were assessed by agarose gel electrophoresis and spectrophotometry. RNA (1µg per triplicate reaction) was reverse transcribed to cDNA using the Taqman Gold Reverse Transcription kit (Applied Biosystems). Real-time RT-PCR (qRT-PCR) was carried out using SYBR^®^ Premix Ex TaqTM, and ALV-J or REV specific primers (Supplementary Table S1). All values were normalized to the endogenous control GAPDH to control for variation. For qRT-PCR of miR-147, we used a miRcute miRNA first-stand cDNA synthesis kit and a miRcute miRNA qPCR detection kit (SYBR Green) (TIANGEN). The miRNA-specific forward primer used for qPCR is 5’-GGTGTGCGGAAATGCTTCTGC-3’. The reverse primer was provided in the miRcute miRNA qPCR detection kit as a primer complementary to the poly (T) adapter. Data were collected on an ABI PRISM 7500 and analyzed via Sequence Detector v1.1 software. All values were normalized to the endogenous control U6 to control for variation. The specific primer of U6 was described in Supplementary Table S1. Assays were performed in triplicate and average threshold cycle (CT) values were used to determine relative concentration differences based on the ΔΔCT method of relative quantization described in the manufacturer’s protocol.

### Cell Counting Kit (CCK-8) for cell proliferation assay

Cell Count Kit-8 was purchased from Solarbio (Beijing, China) and used to examine cell proliferation according to the manufacturer’s instructions.

### ELISA for NF-κB p65 and P-IκBα assay

Chicken NF-κB p65 ELISA kit and Chicken NF-κB P-IκBα ELISA kit were purchased from Senbeijia (Nanjing, China) and used to assay the expression of NF-κB p65 according to the manufacturer’s instructions.

### Statistical analysis

Results are presented as the mean ± standard deviation(s).The T test and ONEWAY ANOVA test was performed using SPSS 13.0 statistical software. A P value less than 0.05 was considered statistically significant.

### Accession numbers

The Illumina small RNA deep sequencing data for the reported miRNAs has been deposited with the NCBI GEO under accession number GSE109105.

### Ethics Statement

The experiment was carried out in strict accordance with the recommendations in the Guide for the Care and Use of Laboratory Animals of the Ministry of Science and Technology of the People’s Republic of China. The protocol was approved by the Committee on the Ethics of Animal Experiments of the Shandong Province (Permit Number: 20150124).

## Acknowledgements

We are grateful to Prof. Yi Tang and Prof. Xiaomin Zhao for their helpful discussion and manuscript revision. We thank Dr Gen Li, Dr Mingjun Zhu and Dr Tianle Chao for their technical assistance.

